# Age-Related Changes in the Rhesus Macaque Eye

**DOI:** 10.1101/2021.07.14.452427

**Authors:** Kira H. Lin, Tu Tran, Soohyun Kim, Sangwan Park, Jiajia Chen, J. Timothy Stout, Rui Chen, Jeffrey Rogers, Glenn Yiu, Sara Thomasy, Ala Moshiri

## Abstract

**Purpose:** To assess age-related changes in the rhesus macaque eye and evaluate them to corresponding human age-related eye disease.

**Methods:** Data from eye exams and imaging tests including intraocular pressure (IOP), lens thickness, axial length, and retinal optical coherence tomography (OCT) images were evaluated from 142 individuals and statistically analyzed for age-related changes. Quantitative autofluorescence (qAF) was measured as was the presence of macular lesions as related to age.

**Results:** Ages of the 142 rhesus macaques ranged from 0.7 to 29 years (mean=16.4 years, stdev=7.5 years). Anterior segment measurements such as IOP, lens thickness, and axial length were acquired. Advanced retinal imaging in the form of optical coherence tomography and qAF were obtained. Quantitative assessments were made and variations by age groups were analyzed to compare with established age-related changes in human eyes. Quantitative analysis of data revealed age-related increase in intraocular pressure, ocular biometry (lens thickness and axial length), and presence of macular lesions. Age-related changes in thicknesses of retinal layers on OCT were observed and quantified. Age was correlated with increased qAF.

**Conclusions:** The rhesus macaque has age-related ocular changes similar to humans. IOP increases with age while retinal ganglion cell layer thickness decreases. Macular lesions develop in some aged animals. Our findings support the concept that rhesus macaques may be useful for the study of important age-related diseases such as glaucoma, macular diseases, and cone disorders, and for development of therapies for these diseases.

## Introduction

There are several human ocular diseases such as cataracts, glaucoma, and macular degeneration that are age-related and vision impairing. Cataracts are the leading cause of blindness globally (Khairallah et al., 2015). Previous models such as mice have been used to study the development of cataracts (Kuck, 1990), however structural differences between the mouse and human eye make the development of cataracts potentially anatomically different from one another. The short lifespan of the mouse also makes it a less ideal model to study potential clinical treatment of cataracts. The second most common cause of blindness globally is glaucoma (Cook and Foster, 2012). Although glaucoma spontaneously occurs in canines, the structural differences between canine and human suggest a different pathological process (Palko et al., 2016). Because of structural differences of the optic nerve head (lamina cribrosa) between primates and rodents, as well as dissimilarities between primate and rodent retinal ganglion cell subtypes, rodent models of glaucoma have significant limitations in the degree to which they are translatable to the human disease. Age related macular degeneration is the largest cause of blindness in the United States. Macular degeneration is a uniquely human disease as most mammals lack complete macular specialization. The non-human primates (NHPs) are the only other mammalian species to have a macula, thus highlighting their requirement as a model for studying macular diseases. Due to the similarities between ocular structures and patterning within primates, it would be beneficial to have primate model systems which replicates age-related changes seen in the human eye.

In order to determine the degree to which these age-related pathologies occur in NHPs, we performed eye exams and imaging tests on rhesus macaques at the Vision Science Laboratory of the California National Primate Research Center (CNPRC) at UC Davis. We examined both eyes of 142 primates of various ages and performed quantitative analysis of the measured data compared to age. Our investigation showed that the rhesus macaque eye undergoes age-related changes corresponding to the most common causes of blindness seen in humans: cataracts, glaucoma, and early stages of macular degeneration.

## Methods

### Animals

All of the animals in this study were rhesus macaques (*Macaca mulatta*) born and maintained at the California National Primate Research Center (CNPRC). The CNPRC is accredited by the Association for Assessment and Accreditation of Laboratory Animal Care (AAALAC) International. Guidelines of the Association for Research in Vision and Ophthalmology Statement for the Use of Animals in Ophthalmic and Vision Research were followed. All aspects of this study were in accordance with the National Institutes of Health (NIH) Guide for the Care and Use of Laboratory Animals. Phenotyping and ophthalmic examinations were performed according to an animal protocol approved by the UC Davis Institutional Animal Care and Use Committee.

### Ophthalmic Data Collection

Phenotypic data were collected from primates previously catalogued for identification of inherited retinal diseases (Moshiri et al., 2019). A small minority of the animals in this study overlapped with a previous study at CNPRC (Yiu et al., 2017). Sedated ophthalmic examination included measurement of intraocular pressure using rebound tonometry (Icare TA01i, Finland), pupillary light reflex testing, external and portable slit lamp examination, as well as dilated (Tropicamide 1%, Phenylephrine 2.5%, Cyclopentolate 1%) indirect ophthalmoscopy. Axial length and lens thickness were measured using a Sonomed Pacscan Plus (Escalon, Wayne, PA, United States). Sedation was achieved by intramuscular injection of ketamine hydrochloride (5-30 mg/kg IM) and dexmedetomidine (0.05-0.075 mg/kg IM). Animals were monitored by a trained technician and a veterinarian at all times.

Color and red-free fundus photographs were obtained with the CF-1 Retinal Camera with a 50° wide angle lens (Canon, Tokyo, Japan). Spectral-domain optical coherence tomography (SD-OCT) with confocal scanning laser ophthalmoscopy (cSLO) was also performed (Spectralis^®^ HRA+OCT, Heidelberg, Germany). A 30-degree horizontal high-resolution raster scan centered on the fovea was obtained using a corneal curvature (K) value of 6.5 mm radius. The Heidelberg eye tracking Automatic Real-Time (ART) software was set at 25 scans for each B-scan acquired. All imaging was done by the same ophthalmic imaging team. All OCT images were taken through the center of the pupil. Speculums were used and corneal hydration was maintained through application of topical lubrication (Genteal artificial tears) approximately every 1-2 minutes during imaging sessions. Focal measurements of each retinal layer thickness were performed using the caliper measurement tool in ImageJ (National Institutes of Health, Bethesda, MD, United States) on a horizontal line scan through the foveal center. The foveal center was determined on optical coherence tomographic images where the fovea had the greatest depth. The manufacturer’s scale on the OCT image was used to calibrate ImageJ prior to taking measurements of each layer. Each retinal layer thickness was measured at 1.5mm temporal to the foveal center, at the foveal center, and 1.5mm nasal to the foveal center. The layers measured included the nerve fiber layer (NFL), ganglion cell layer (GCL), inner plexiform layer (IPL), inner nuclear layer (INL), outer plexiform layer (OPL), outer nuclear layer (ONL), photoreceptor inner segments (IS), photoreceptor outer segments (POS), retinal pigmented epithelium (RPE), choriocapillaris (CC), outer choroid (OC), and total retinal thickness (TRT).

The thickness of each layer was segmented manually and measured boundary to boundary (Litts et al., 2018; Staurenghi et al., 2014). As reported previously, the choroidal-scleral junction cannot be reliably visualized in every animal, which may confound OC measurements (Yiu et al. 2017). Averages were calculated for each layer at each location for each eye of each individual primate.

The Spectralis device was also used to obtain blue-peak fundus autofluorescence in a quantitative fashion. After SD-OCT images were captured, the device was turned to qAF mode to capture 30° x 30° qAF images. The device was calibrated with an internal master fluorescence reference and set to excitation light of 488 nm and a long-pass barrier filter starting at 500 nm increasing to 80 nm. The retina was exposed to 488 nm blue excitation light for 30 seconds to achieve photobleaching. Intensity was adjusted for using an internal fluorescence reference to enable quantification of autofluorescence (AF) and normalizing AF units. Images were captured from the central macula with minor variations in laser power and detector sensitivity in between imaging sessions. Each eye had three series of 12 rapid succession images acquired and the mean of the three sequences were calculated using the manufacter’s qAF software. We occasionally encountered opacities in ocular media that limited the quality of imaging. Quantitative autofluorescence (qAF) images were chosen based on grading between two individuals (K.H.L. and T.M.T.). Images were graded from 0-3 (0 being ineffectual and 3 being high quality) with only images of a grade of 2 or above being evaluated. Acceptable images were measured using a Delori grid centered on the fovea and expanded to the tangential edge of the optic nerve. The main measure for each image is the qAF8, which was acquired after selecting the middle eight segments to exclude vessels and other noise and calculating the mean value. The mean qAF8 value was calculated as the mean of the three images per eye and averaged between the two graders.

### Histology

The enucleated eye was dissected to isolate the macula and immediately fixed using freeze substitution and embedded in paraffin as previously described (Sun et al., 2015). The macular region was sectioned at 5 μm on a microtome and sections collected on slides and dried. Using standard protocols, sections were treated with cold acetone, hematoxylin, 0.5% HCl in 70% ethanol, eosin, 95% and 100% ethanol and Histo-Clear for light microscopic examination.

### Statistical Analysis

Descriptive statistics were used to evaluate age-related changes. Panel regression was used to determine statistical significance (*P* < 0.05) treating each eye as the unit of analysis linked by the primate to account for within-subject correlation, and each primate’s panel consisted of right eye and left eye measurements when applicable. Analyses were performed in Microsoft Excel (Microsoft Corporation, Redmond, WA, United States) and STATA 16 (StataCorp, College Station, TX, United States).

## Results

Data were collected from a total of 142 individuals. Ages of the rhesus macaques ranged from 0.7 to 29 years (mean = 16.4 years, stdev = 7.5 years). The age distribution is shown in Table 1.

**Table 1.**
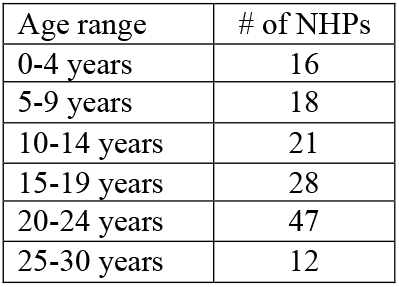
Age distribution of rhesus macaques (n = 142) by age. The mean was 16.4 years (stdev = 7.5 years). Median was 19.2 years.

| Age range | # of NHPs |
| --- | --- |
| 0-4 years | 16 |
| 5-9 years | 18 |
| 10-14 years | 21 |
| 15-19 years | 28 |
| 20-24 years | 47 |
| 25-30 years | 12 |

**Table 2.** Age-related changes in retinal layers as measured on OCT. Thickness measurements in other layers did not show age-related changes. Abbreviations: NFL: nerve fiber layer. GCL: ganglion cell layer. IPL: inner plexiform layer. INL: inner nuclear layer. OPL: outer plexiform layer. ONL: outer nuclear layer. IS: inner segments. POS: photoreceptor outer segments. RPE: retinal pigmented epithelium. CC: choriocapillaris. OC: outer choroid. TRT: total retinal thickness.

| Retinal Layer | Slope Coefficient ($\mu\text{m}$ per increase in year of age) | P-value | Correlation Coefficient |
| --- | --- | --- | --- |
| Nasal GCL | -0.30 | 0.019 | 0.56 |
| Temporal GCL | -0.37 | 0.001 | 0.74 |
| Temporal INL | -0.19 | 0.025 | 0.78 |
| Nasal POS | 0.16 | 0.003 | 0.81 |
| Temporal POS | 0.26 | <0.001 | 0.76 |
| Foveal POS | 0.36 | <0.001 | 0.51 |
| Nasal CC | 0.34 | <0.001 | 0.56 |
| Temporal CC | 0.38 | <0.001 | 0.56 |
| Foveal CC | 0.27 | <0.001 | 0.75 |
| Nasal OC | 3.66 | <0.001 | 0.28 |
| Temporal OC | 3.20 | <0.001 | 0.74 |
| Foveal OC | 3.58 | <0.001 | 0.60 |

Intraocular pressure (IOP) measurements were collected from a total of 142 individuals, 284 eyes. We found that IOP increased (Figure 1) by 0.165 mm Hg per each year of age (*P* < 0.001). Lens thickness and axial length measurements were collected from 114 individuals, 228 eyes total. Lens thickness measurements suggested a positive correlation with age, with an increase of 7.3 ± 5.2 μm per year in age (*P* = 0.162, Figure 2A). Axial length increased by 52.8 ± 11.3 μm per year of age (*P* < 0.001, Figure 2B).

**Figure 1.**
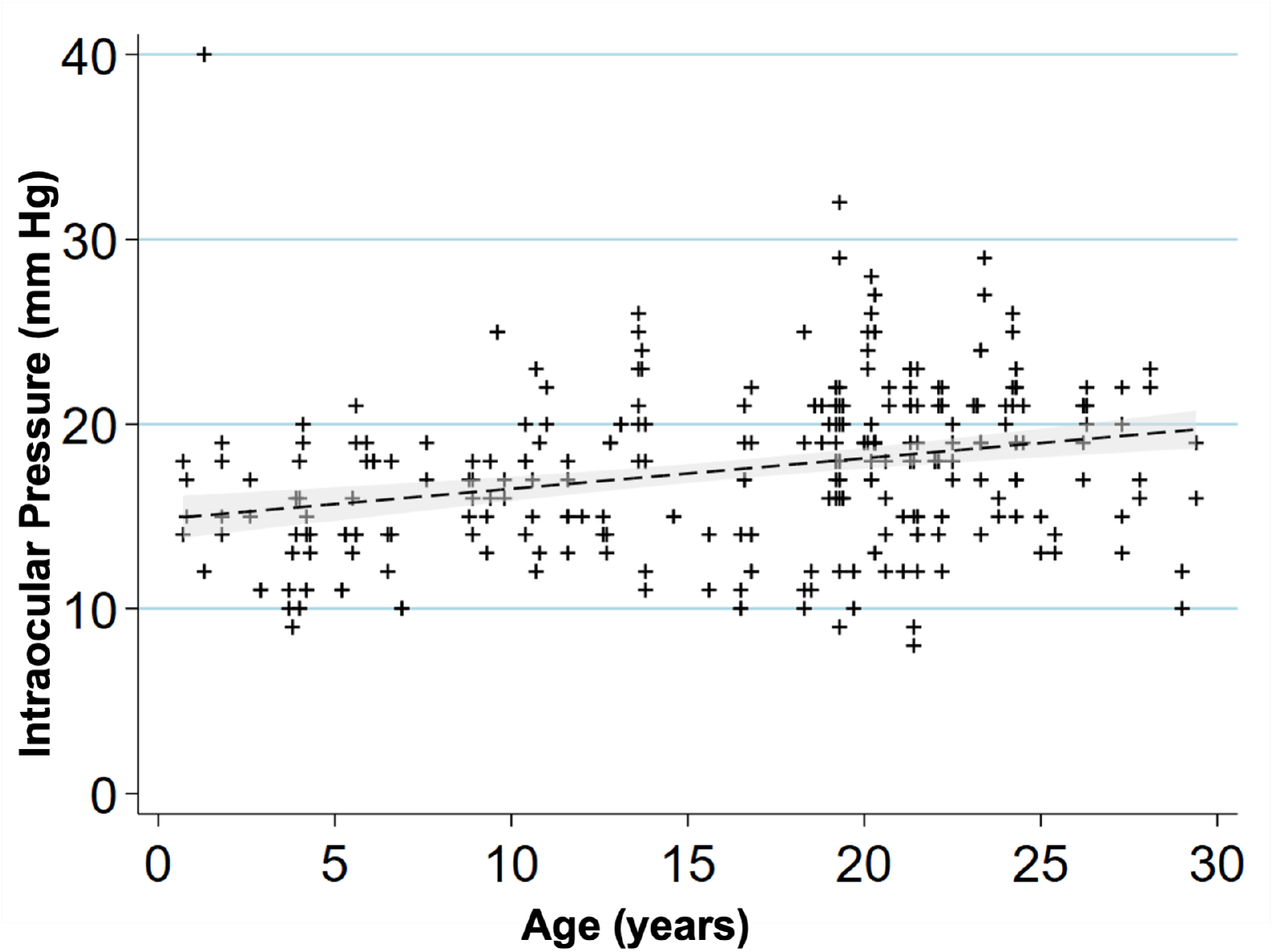
Intraocular pressure increases with age in rhesus macaques. Scatterplot showing relationship of age (years) and intraocular pressure (mm Hg) for 142 macaques, 284 eyes. IOP increased by 0.165 mm Hg per increase in year of age (*P* < 0.001). A linear regression line is fitted with a 95% confidence interval in the grey area.

**Figure 2.**
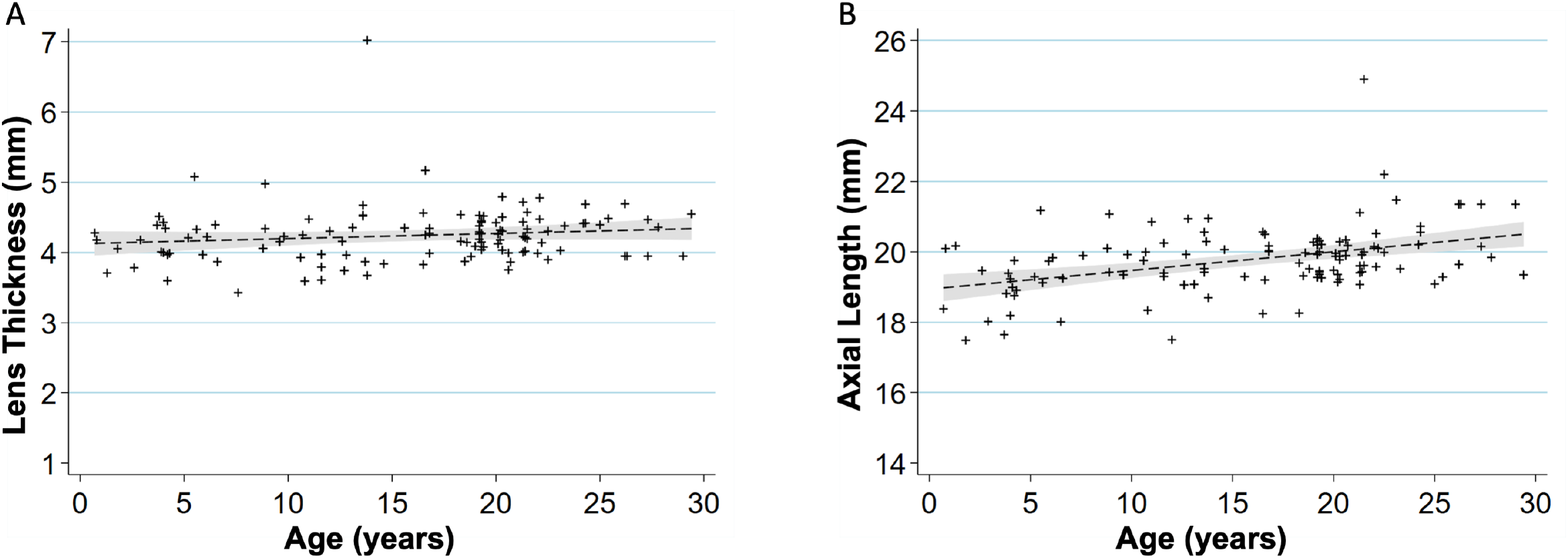
Age-related changes in ocular biometry. **(A)** Scatterplot showing relationship between age (years) and lens thickness (mm) (n = 114 primates, 228 eyes). The lens thickness increases 7.3 μm per increase in year of age (*P* = 0.162). **(B)** Scatterplot showing relationship between age (years) and axial length (mm) (n = 114 primates, 228 eyes). The axial length increases 52.8 μm per increase in year of age (*P* < 0.001). A linear regression line that has been fitted with a 95% confidence interval is shown in the grey area of each graph.

### Retinal Findings

Fundus imaging was evaluated on 78 individuals (77 right eyes, 78 left eyes). Observations by indirect ophthalmoscopy revealed yellow-white punctate macular lesions of likely lipoidal degeneration (Anderson et al., 2006; Curcio et al., 2010; Rudolf et al., 2019; Yiu et al., 2017; Yiu et al., 2020) present in 66 individuals (age range = 4.2-29.4 years, mean age = 17.8 years, median age = 19.3 years, 47% of total study subjects). These lesions, reminiscent of small hard drusen in humans, were identifiable on fundus imaging (Figure 3A-D & 4) and were distinct from soft drusen-like macular lesions (Figure 3E) observed in a smaller subset of animals. Hematoxylin and eosin stain of a histological section of a rhesus macaque retina clinically graded with extensive punctate macular lesions shows numerous translucent spheroidal lesions on the apical aspect of the retinal pigmented epithelium cells, near the junction of the photoreceptor outer segments (Figure 3F). Animals with extensive punctate macular lesions had granular hyperreflective foci seen in the RPE band on OCT. Autofluorescence imaging, fundus photos, and an OCT scan through the macular lesions are shown in Figure 4, in contrast to that seen in animals with no macular lesions (Supplemental Figure 1). There were 74 individuals without punctate macular lesions (age range = 0.7-27.8 years, mean age = 14.6 years, median age = 18.8 years). The age distributions of individuals with and without punctate macular lesions are shown in Figure 5A. Forty-two individuals had punctate macular lesions visible by indirect ophthalmoscopy and on fundus images. The mean age of the animals with punctate macular lesions (17.8 ± 6.2 years) was significantly older than the mean age of animals without them (14.6 ± 8.1 years), suggesting punctate macular lesions are an age-related phenomenon (*P* = 0.009) (Anderson et al., 2006; Curcio et al., 2010; Yiu et al., 2017; Yiu et al., 2020). Further grouping of individuals with various degrees (clinically graded by indirect ophthalmoscopy as few, moderate, or extensive) of punctate macular lesions is shown in Figure 5B. While the presence of punctate macular lesions may be age-related, the extensiveness of punctate macular lesions may not correlate solely with age.

**Figure 3.**
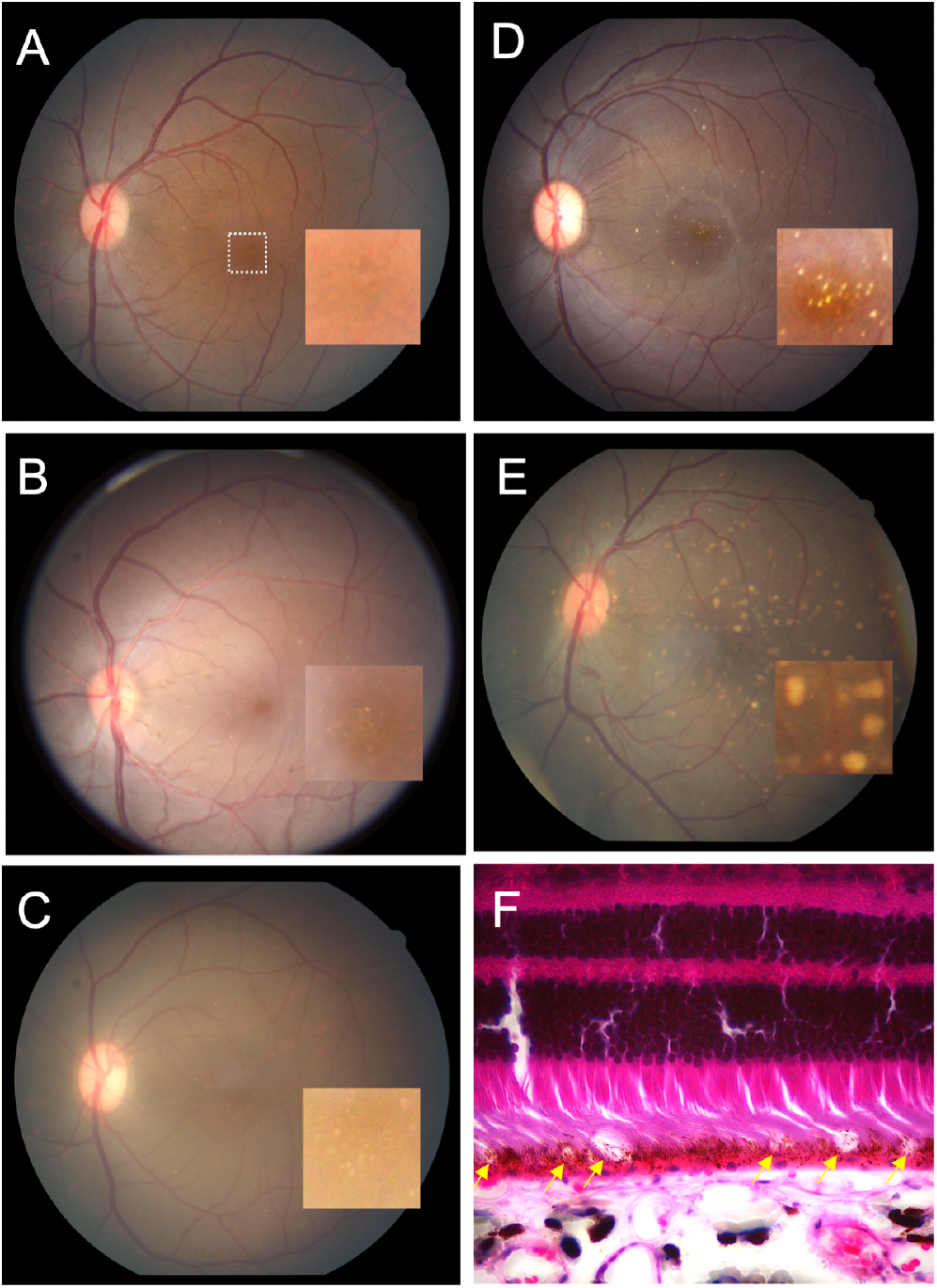
Macular lesions in rhesus macaques. Grading of macular lesions by clinical examination based on number and area of the posterior pole involved. Inset in A-E is a magnification of the foveal center (dashed box in A) highlighting the lesions. **(A)** Color fundus photo Group 0: no punctate macular lesions. **(B)** Color fundus photo Group 1: few punctate macular lesions, typically limited to the foveal avascular zone. **(C)** Color fundus photo Group 2: moderate punctate macular lesions, typically in the fovea with a few outside the foveal avascular zone. **(D)** Color fundus photo Group 3: extensive punctate macular lesions, typically in the fovea and throughout the macula. **(E)** Color fundus photo Group 4: soft drusen-like macular lesions. **(F)** Hematoxylin and eosin stain of a histological section of a rhesus macaque retina clinically graded with extensive punctate macular lesions. Numerous translucent spheroidal lesions in the outer retina in the region of the photoreceptor outer segments and the retinal pigmented epithelium are seen (yellow arrows). The section is from the macular region as evidenced by multiple rows of nuclei in the retinal ganglion cell layer.

**Figure 4.**
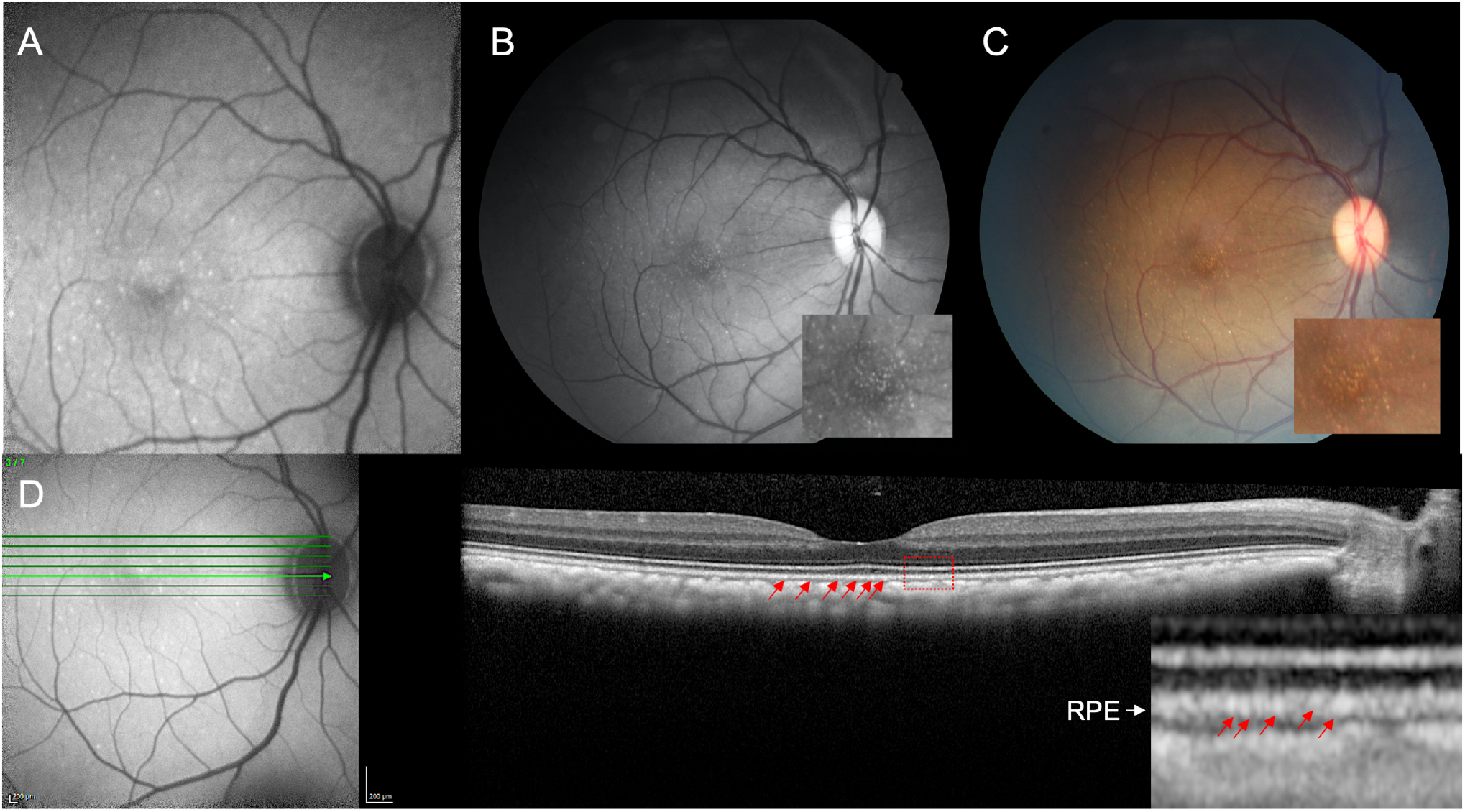
Multimodal retinal imaging of punctate macular lesions. Imaging of punctate macular lesions by (A) fundus autofluorescence, (B) red-free and (C) color fundus imaging in an animal clinically graded as extensive. Enlarged inset images show lesions in the fovea. Optical coherence tomography (D) shows punctate hyperreflectivity at the level of the RPE (red arrows) in the enlarged inset demarcated by the red box.

**Figure 5.**
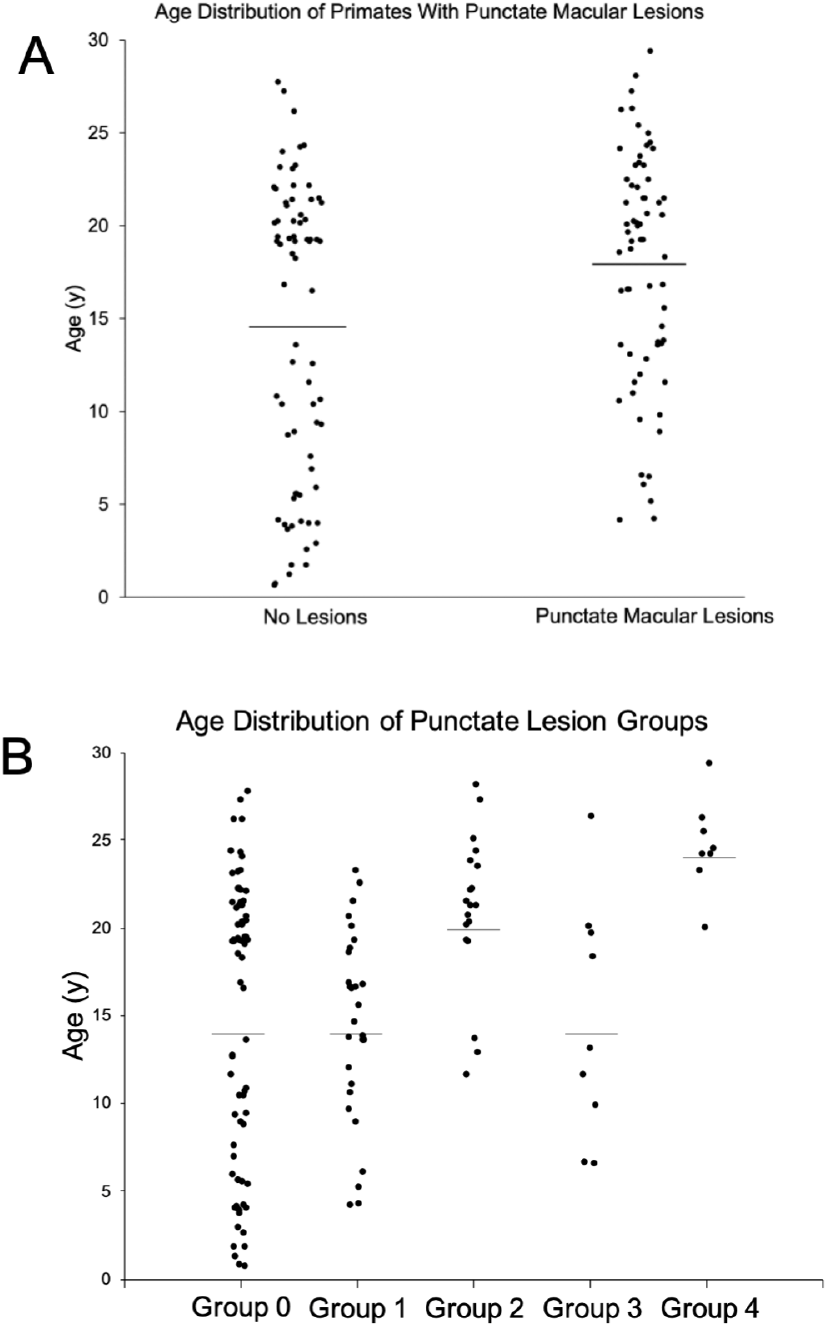
Macular lesions and age distribution. **(A)** Age distribution of rhesus macaques with and without punctate macular lesions (*P* = 0.009). No punctate macular lesions (n = 74, age range = 0.7-27.8 years, mean age = 14.6 years, median age = 18.8 years). Punctate macular lesions (n = 66, age range = 4.2-29.4 years, mean age = 17.8 years, median age = 19.3 years). Macaques with punctate macular lesions = 47% of total study primates. Line represents mean age. **(B)** Age distribution of rhesus macaques with various levels of punctate macular lesions. Group 0: no punctate lesions (n = 74, age range = 0.7-29.0 years, mean age = 14.6 years, median age = 19.8 years). Group 1: few lesions (n = 30, age range = 4.2-23.3 years, mean age = 14.9 years, median age = 16.0 years). Group 2: moderate lesions (n = 19, age range = 11.6-28.1 years, mean age = 20.9 years, median age = 21.3 years). Group 3: extensive lesions (n = 9, age range = 6.5-26.3 years, mean age = 14.7 years, median age = 13.1 years). Group 4: soft drusen-like macular lesions (n = 8, age range = 20.0-29.4 years, mean age = 24.6 years, median age = 24.3 years). Line represents mean age.

### Optical Coherence Tomography Segmentation

Tomographic images of the macula were evaluated on 60 individuals, 49 right eyes and

49 left eyes. To determine if there are structural changes in the retina over time, individual retinal layer thicknesses were plotted against age (Figures 6 & 7). Significant changes in layer thickness with increase in age were found in the retinal ganglion cell layer (GCL), inner nuclear layer (INL), photoreceptor outer segments (POS), choriocapillaris (CC), and outer choroid (OC) consisting of both the Sattler and Haller layers of the choroid. Significant reduction in thickness was found in the GCL both nasal (*P* = 0.019) and temporal (*P* = 0.001) to the fovea. The nasal side decreased by 0.30 μm and the temporal side by 0.37 μm for each year of age. The thickness of the INL reduced only temporal to the foveal by 0.19 μm per year of age (*P* = 0.025). The POS increased in thickness on the nasal side (0.16 μm per year, *P* = 0.003), temporal side (0.26 μm per year, *P* < 0.001), and foveal center (0.36 μm per year, *P* < 0.001). The CC thickness increased in the nasal side (0.34 μm per year, *P* < 0.001), temporal side (0.38 μm per year, *P* < 0.001), and foveal center (0.27 μm per year, *P* < 0.001). Finally, the OC thickness increased in the nasal side (3.66 μm per year, *P* < 0.001), temporal side (3.20 μm per year, *P* < 0.001), and foveal center (3.58 μm per year, *P* < 0.001). The other layers were not found to vary significantly with age.

**Figure 6.**
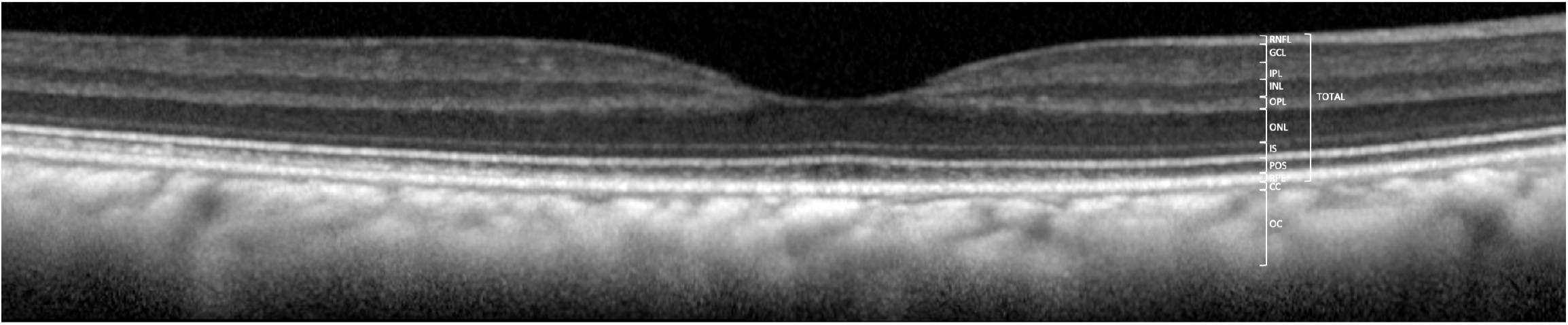
Optical Coherence Tomography (OCT) images and measurements. **(A)** Measurement of retinal and choroidal layers (n = 60 primates, 98 eyes). All retinal layers were measured at 1.5 mm on both temporal and nasal sides and in the foveal center. Total retinal thickness is measured from the nerve fiber layer (NFL) to the retinal pigment epithelium (RPE). Total retinal thickness caliper is shown at 1.5 mm eccentricity.

**Figure 7.**
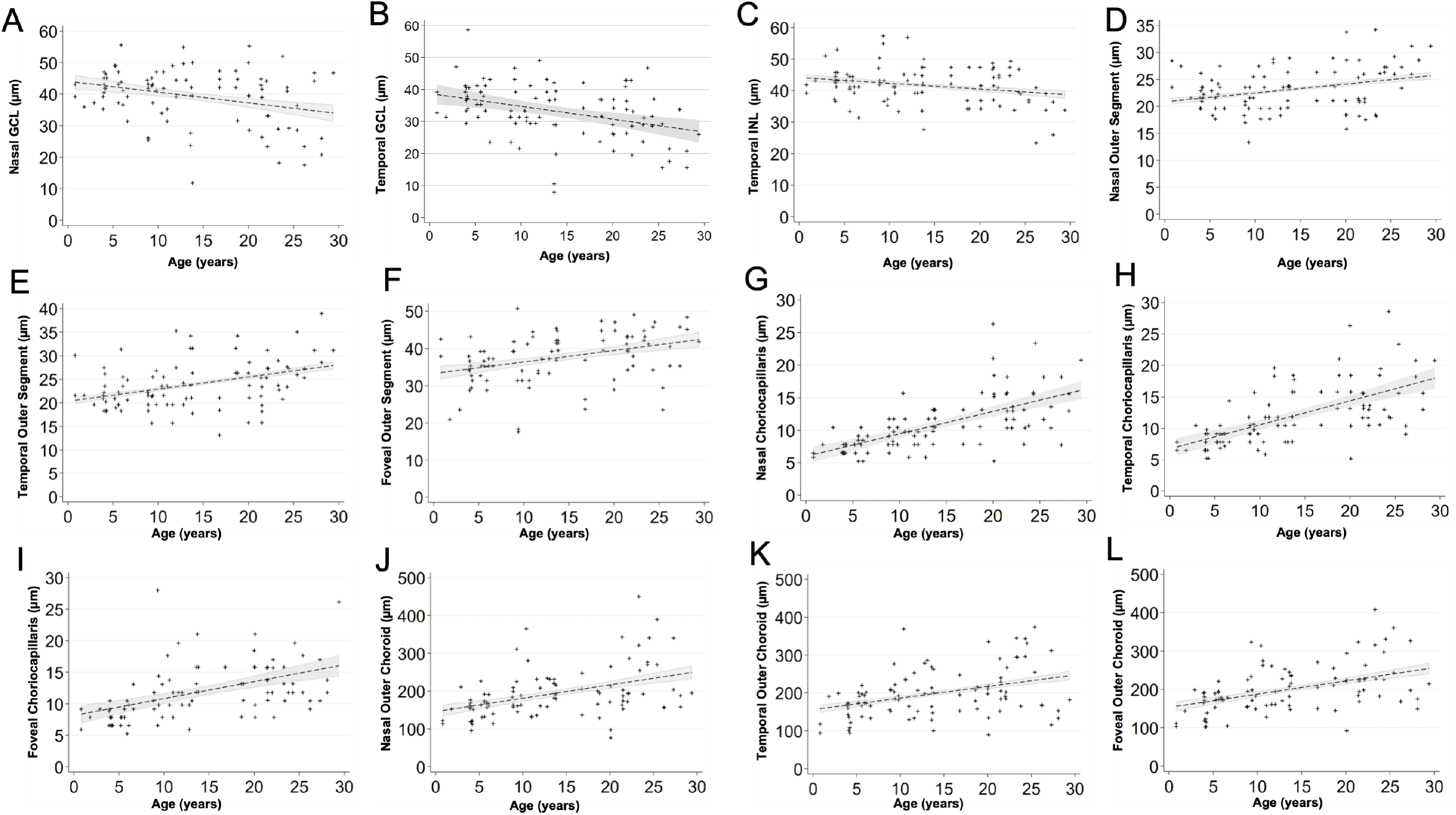
Scatterplots show the relationship between age (years) and the thickness of each layer (μm). **(A)** Nasal GCL decreased by a factor of 0.30 μm per increase in year of age (*P* = 0.019). **(B)** Temporal GCL decreased by a factor of 0.37 μm per increase in year of age (*P* = 0.001). **(C)** Temporal INL decreased by a factor of 0.19 μm per increase in year of age (*P* = 0.025). **(D)** Nasal POS increased by a factor of 0.16 μm per increase in year of age (*P* = 0.003). **(E)** Temporal POS increased by a factor of 0.26 μm per increase in year of age (*P* < 0.001). **(F)** Foveal POS increased by a factor of 0.36 μm per increase in year of age (*P* < 0.001). **(G)** Nasal CC increased by a factor of 0.34 μm per increase in year of age (*P* < 0.001). **(H)** Temporal CC increased by a factor of 0.38 μm per increase in year of age (*P* < 0.001). **(I)** Foveal CC increased by a factor of 0.27 μm per increase in year of age (*P* < 0.001). **(J)** Nasal OC increased by a factor of 3.66 μm per increase in year of age (*P* < 0.001). **(K)** Temporal OC increased by a factor of 3.20 μm per increase in year of age (*P* < 0.001). **(L)** Foveal OC increased by a factor of 3.58 μm per increase in year of age (*P* < 0.001).

### Quantitative Autofluorescence (qAF) analysis

The qAF8 data of 66 eyes of 44 individuals was evaluated. An image of the Delori grid in Heidelberg’s qAF mode is shown in Figure 8A. qAF8 consisted of the 8 segments of the middle ring of the pattern computed to a mean (Gliem et al.,2016). We observed an increase in qAF8 by a factor of 1.021 units per year of age (*P* = 0.006, Figure 8B).

**Figure 8.**
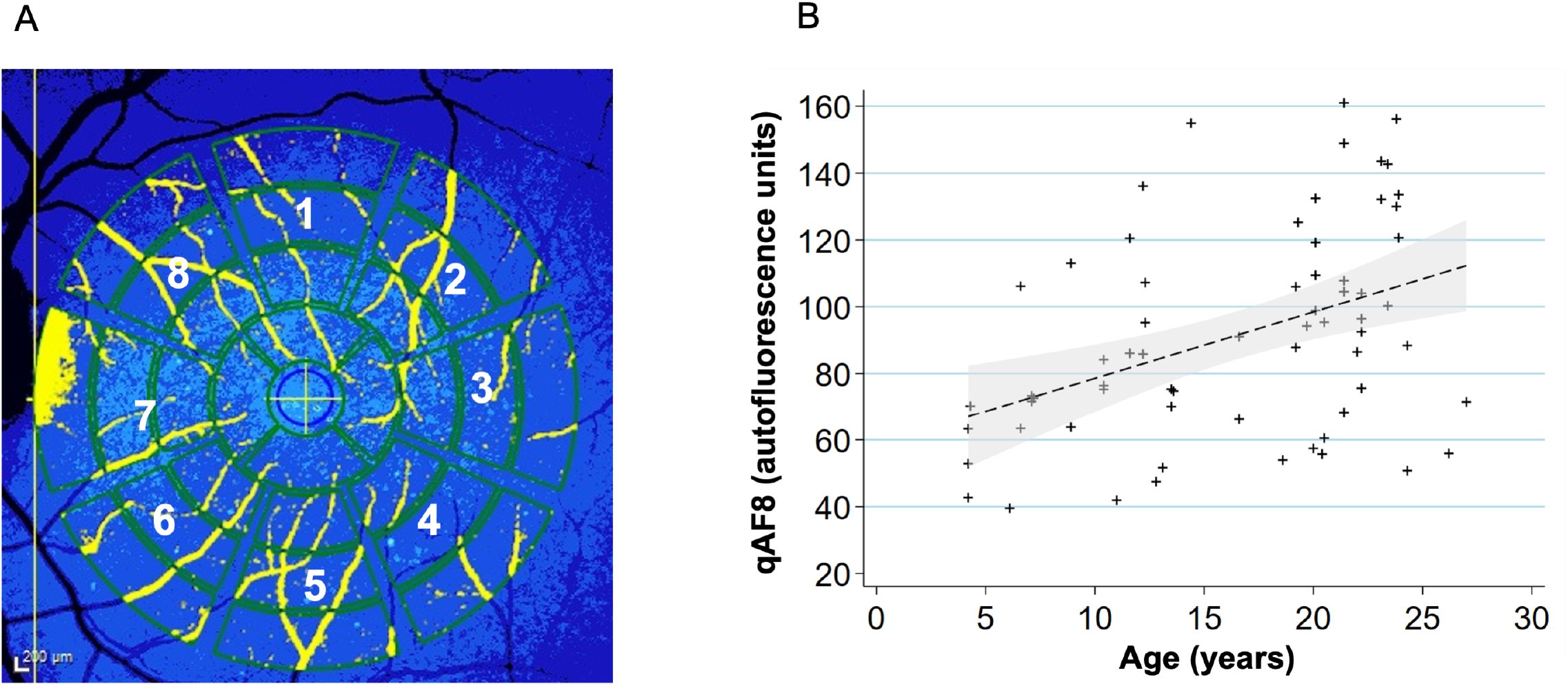
Quantitative autofluorescence analysis (n = 44 primates, 66 eyes) **(A)** To acquire qAF8, the eight middle segments (numbered octants) of the perifoveal Delori grid was used in Heidelberg’s qAF mode. The center circle is placed over the foveal center and the grid is expanded until it touches the tangential edge of the optic nerve. Vessels are automatically excluded and the qAF measurements in each octant are normalized to the internal autofluorescence standard shown in the blue bar at the top of the image. **(B)** Scatterplot showing relationship between qAF8 and age (years). Mean (standard deviation): 91.4 (31.3). The mean qAF8 increased by a factor of 1.021 autofluorescence units per increase in year of age (*P* = 0.006). A linear regression line has been fitted with a 95% confidence interval in the grey area.

## Discussion

### Age-related changes in Intraocular Pressure (IOP), Lens Thickness, and Axial Length

It is believed that rhesus macaques age about 3 times faster than humans (Simmons, 2016). They generally reach adulthood (sexual maturity) by 5 years, are considered geriatric after 20 years, and can live up to 30 years. In our cohort of macaques, we observed a significant increase in IOP with age (*P* < 0.005). A similar trend in humans has been reported across ethnicity (Wong et al., 2009; Klein et al., 1992). Consistent with prior reports, we also observed a positive trend of the proportion of primates that had an IOP >21 mm Hg. One primate in age-group 5-9 years old (6%), 4 primates in age-group 10-14 years old (19%), 5 primates in age-group 15-19 years old (18%), and 16 primates in age-group 20-24 years old (34%) were observed to have an IOP >21 mm Hg in at least one of their eyes. Age-group 25-30 years old had 3 primates (25%) that had an IOP >21 mm Hg, but given the advanced age of these primates this group was small. This age-related IOP elevation suggests that rhesus macaques may develop corresponding optic nerve damage similar to primary open angle glaucoma. Furthermore, we did find thinning of the retinal ganglion cell layer with age in this study. Dawson et al. (1993) also found evidence of primary open angle glaucoma within a closed colony of rhesus monkeys on the island of Cayo Santiago. Thirty-one individuals had elevated intraocular pressures of ≥ 21 mm Hg. However, they did not find any age-related significance within the affected individuals. A normative primate and ocular hypertensive primate were examined in greater depth and the hypertensive individual showed decreased optic nerve fibers and reduced axon densities at various sites.

It has been shown that the fully developed normal human lens is 4.0 mm in thickness at age 20, but increases to 4.3 mm at the age 40 of years, 4.45 mm at 50 years, 4.7 mm at 60 years, and above 4.7 mm after 60 years of age (Levin et al., 2011). The rate of increase in crystalline lens thickness has been estimated at 0.15-0.20 mm per decade of life (Roters et al., 2002). The macaque lens is 4.24 ± 0.41 mm in thickness with an average age of 15.7 years. A previous study in a smaller cohort of rhesus monkeys found a significant age-related increase of 0.03±0.01 mm per year (*P* = 0.002) in lens thickness (Wendt et al., 2008). However, lens thickening with age did not meet statistical significance in our cohort (*P* = 0.162).

The average axial length measurement of the rhesus eye was determined to be 19.77 ± 0.97 mm with a significant age-related increase of 52.8 μm per year (*P* < 0.001). The average axial length for animals aged 0-4 was 18.89 ± 0.83 mm while the animals age 25-30 measured 20.28 ± 0.97 mm. In humans, the newborn eye is 16 mm and continues to grow to approximately 24 mm (Goldschmidt, 1969). In humans, age-related axial length increase causes myopia, and this process can lead to pathological consequences and vision loss (Flores-Moreno et al., 2013; Mutti et al., 2007; Ohno-Matsui, 2016; Silva, 2012). We did observe rare examples of high myopia in our cohort of monkeys and among normal eyes axial length did increase similar to what has been reported in the clinical literature. Therefore, rhesus monkeys may serve as reasonable models for the study of normal axial length increase and potentially also for pathologic myopia (Smith et al., 2012).

### Age-related changes on Retinal OCT and qAF

Drusen are lesions characteristic of age-related macular degeneration (AMD). Our cohort of rhesus macaques had punctate macular lesions, which appear to be an age-related phenomenon as the group of animals with them were significantly older than those without them. However, they could occur even in young animals and the extensiveness of these lesions did not correlate with age, suggesting genetic or environmental factors may also be involved. Punctate macular lesions seem to correspond to vacuolated spherules at the apical aspect of the RPE cells on histopathology, and animals with these lesions have granular hyperreflectivity of the RPE band on OCT. Rudolf et al. reported essentially identical fundus lesions which corresponded histologically to lipoidal degeneration of individual discrete RPE cells. These lesions had no discernible RPE band alteration on OCT, perhaps due to differences in image acquisition. The punctate lesions in our cohort likely represent the same biological process. Soft drusen-like lesions more reminiscent of those seen in humans with AMD were also observed, but only in a smaller cohort of geriatric animals. The presence of drusen in humans is associated with age and their development is similar to other age-related disease processes (Anderson et al., 2002). As soft drusen-like lesions were only observed in geriatric animals, we believe they may represent early AMD in rhesus macaques. Examples of advanced AMD, geographic atrophy or choroidal neovascularization, were not seen in this study. The lack of geographic atrophy or choroidal neovascularization in this study is consistent with these forms of advanced AMD not having been seen in any of the published surveys of macaque eyes, except in the context of pathologic myopia (Stafford et al. 1984).

We found significant decreases in layer thickness with age in the GCL and also in the temporal INL. Significant increases in thickness were found in the POS, CC, and OC. Demirkaya et al. (2013) showed that humans also have a significant decrease in the pericentral GCL and peripheral INL. However, their study also found a decrease in foveal POS layer thickness contrary to our data. Furthermore, Patel et al. (2014) found significant age-related thinning of the NFL layer, while our cohort showed no significant changes in this layer with age. The choroidal thickening with age in this study is at odds with human aging data (Ramrattan et al., 1994; Ruiz-Medrano et al., 2017) and may be due to the difficulty in measuring the choroidal-scleral junction in NHPs. The discrepancies could be due to true species differences, or secondary to a broader range of subjects in our study. The different methods of measuring layer thickness may also play a role as both human studies used an automated measuring program while our layers were measured manually. Variability in the precise definitions of retinal layer boundaries may also play a role.

In rhesus macaques, qAF was significantly lower than in humans with the average of 91.4 qAF units in rhesus macaques compared to a human mean of 253.6-283.9 qAF units (Greenberg et al., 2013; Wang et al., 2019). We did observe an increase in qAF with increasing age consistent with human data (Armenti et al., 2016; Greenberg et al., 2013; Wang et al., 2019). However, we did not observe a continuous decline in qAF units, which occurs at age 75 years in humans (Table 3, Armenti et al., 2015). The higher qAF in humans may be due to higher absolute age in humans which may allow for decades of accumulation of lipofuscin.

**Table 3.** Mean qAF units across age (n=44 primates, 66 eyes). The mean age was 17.1 years (stdev = 7.1 years). Median was 19.6 years. The mean qAF was 91.4 units (stdev = 31.3 units). Median was 88.1 units.

| Age range | Mean qAF Units |
| --- | --- |
| 0-4 years | 81.1 |
| 5-9 years | 94.9 |
| 10-14 years | 84.1 |
| 15-19 years | 112.8 |
| 20-24 years | 89.7 |
| 25-30 years | 99.3 |

Alternatively, visual cycle metabolism in the outer retina, levels of melanin in the choroidal layer, or other species-specific differences may be responsible for these qAF observations.

The age-related changes in the rhesus macaque eye show consistent similarities to its human counterpart. Age-related changes documented in humans such as IOP elevation, axial length increase, presence of punctate macular lesions, soft drusen-like lesions, and increasing qAF were confirmed in the eye of rhesus macaques. Our findings support the use of the NHP eye as a model for advanced translational vision science research, especially those related to macular and cone-disorders and age-related eye diseases.

## Acknowledgements

The authors thank Monica Motta and Michelle Ferneding for expertise in ophthalmic imaging, electrophysiology, data management, and research support.

---

**Supplemental Figure 1.** Fundus without Macular Lesions

**(A)** Autofluorescence Image

**(B)** Red-free Fundus Photo

**(C)** Color Fundus Photo

**(D)** OCT Scan

