## Supplementary figures and images for "Age-Related Changes in the Rhesus Macaque Eye"

### Supplemental Figure 1

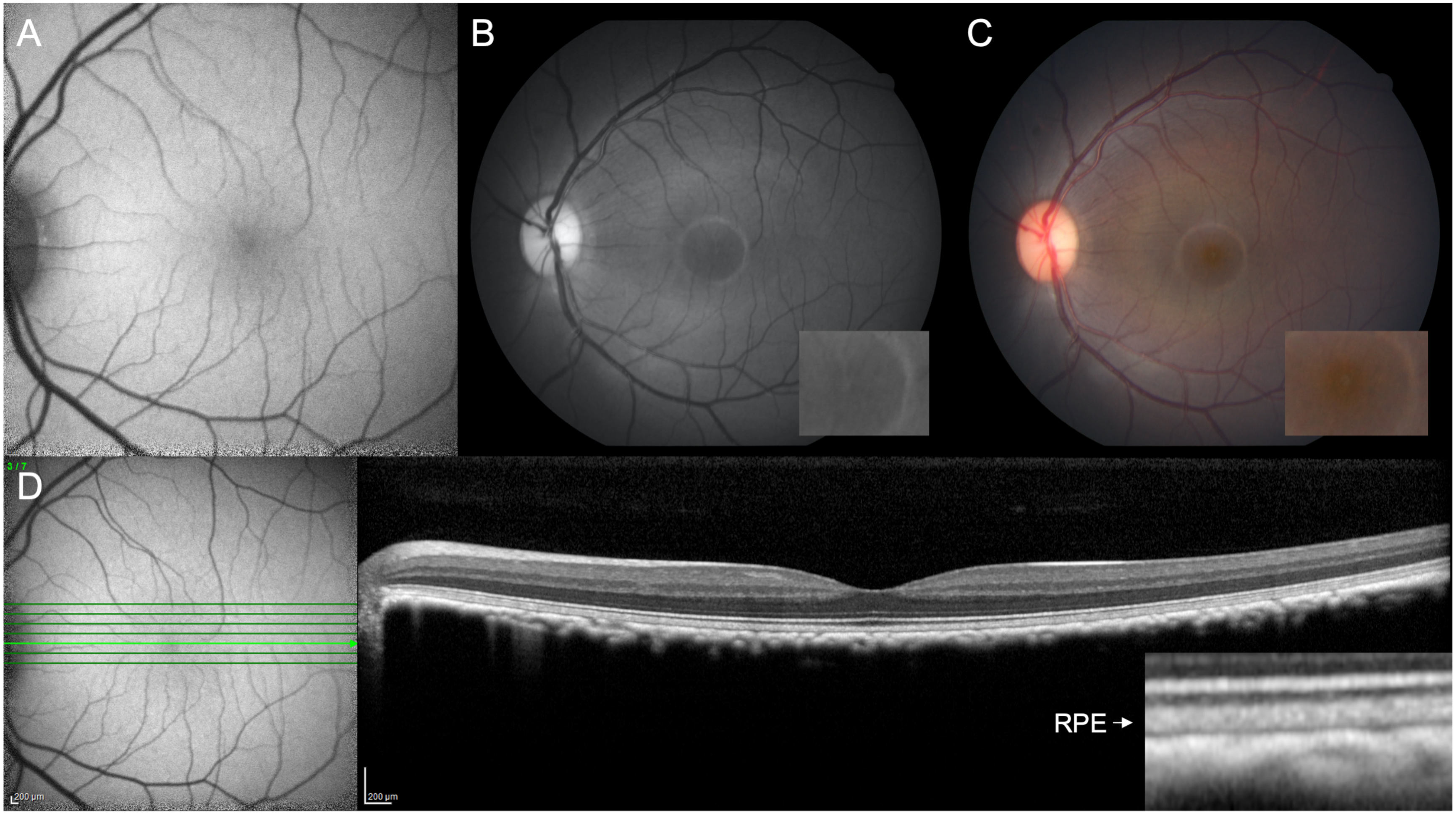
